# Up-regulation of Retrograde Response in yeast increases glycerol and reduces ethanol during wine fermentation

**DOI:** 10.1101/2023.12.01.568406

**Authors:** Víctor Garrigós, Beatriz Vallejo, Esperanza Mollà-Martí, Cecilia Picazo, Emilien Peltier, Philippe Marullo, Emilia Matallana, Agustín Aranda

## Abstract

Nutrient signaling pathways play a pivotal role in regulating the balance between metabolism, growth and stress response depending on the available food supply. They are key factor for the biotechnological success of the yeast *Saccharomyces cerevisiae* during food-producing fermentations. One such pathway is the Retrograde Response, which controls the use of carbon backbones for the synthesis of amino acids derived from alpha-ketoglutarate, such as glutamate and lysine. The repressor *MKS1* is linked to the TORC1 complex and negatively regulates this pathway. Deletion of *MKS1* on a variety of industrial strains causes an increase in glycerol in winemaking, brewing and baking conditions. This increase is accompanied by a reduction in ethanol production in grape juice fermentations in four commercial wine strains. Interestingly, this does not lead to an increase in volatile acidity, as the levels of acetic acid actually decrease. Aeration during winemaking usually increases acetic acid levels, but this effect is reduced in the *MKS1* mutant. Although there is an improvement in the metabolites of oenological interest, it comes at a cost, as the mutant showed slower fermentation kinetics when grown in grape juice, malt, and laboratory media using glucose, sucrose, and maltose as carbon sources. The deletion of *RTG2*, an activator of the Retrograde Response that acts as an antagonist of *MKS1*, also results in a defect in wine fermentation speed. These findings suggest that the deregulation of this pathway causes a fitness defect. Therefore, manipulating the repressor *MKS1* is a promising approach to modulate yeast metabolism and produce low-ethanol drinks.

## Introduction

Food-producing yeasts require a rich and balanced growth media for optimal performance. Insufficient or unbalanced nutrients can cause stress in biotechnological conditions, leading to cell growth arrest and delayed fermentations (Orozco et al. 2019). The mechanisms that respond to nutrient abundance and starvation are called nutrient signaling pathways, and they have been well characterized in laboratory conditions for the budding yeast *Saccharomyces cerevisiae* (Conrad et al. 2014). Those pathways sense the levels of each nutrient to promote or cease cell growth and stress response in a coordinated way. The presence of glucose induces cAMP synthesis and Protein Kinase A activation, while abundance of nitrogen activates TORC1 complex to promote protein synthesis. Both pathways are coordinated and share common targets in order to achieve a subtle control of metabolism. Knowledge of such pathways in laboratory conditions are useful to understand the behavior of industrial yeast, but differences in growth media and conditions must be taken into account. A phenomic analysis of a wine strains carrying deletions in key genes of different signaling pathways pointed to the relevance of PKA in wine fermentation and the relationships between carbon and nitrogen repression mechanisms (Vallejo et al. 2020b). However, behavior of such pathways cannot be taken for granted in industrial environments, as an early induction during grape juice fermentation of markers that are usually linked with carbon and nitrogen starvation when there is no nutrient shortage was shown (Vallejo et al. 2020a).

Retrograde Response (RR) includes a subset of genes that are coordinately expressed and that are involved in the biosynthesis of amino acids from TCA intermediate α-ketoglutarate (Jazwinski 2013; da Cunha et al. 2015). This is a main node for amino acid synthesis as it is used by glutamate dehydrogenase to give glutamic acid, being the main point for ammonium incorporation. RR does not control all mitochondrial TCA enzymes, just the ones from oxaloacetate to α-ketoglutarate, plus other enzymes that produce precursors in the cytoplasm and peroxisomes, like cytosolic pyruvate carboxylase *PYC1* and peroxisomal citrate synthase *CIT2*. Their transcriptional activation happens through transcription factors Rtg1/3 when mitochondrial function is low or glutamic acid is scarce, Upstream regulation in dependent on an activator, Rtg2, and a repressor, Mks1, that interact to each other. Mks1 is described as a pleiotropic negative transcriptional regulator as it was originally described as negative regulator in Ras-cAMP signaling (Matsuura and Anraku 1993). Later it was clearly identified as a TORC1 complex target (Dilova et al. 2002), so it clearly coordinates different signals to better control amino acid metabolism. Transcriptomic analysis showed a tight control of lysine biosynthetic genes by *MKS1*,as most enzymes in the pathway from α-ketoglutarate into lysine are upregulated after its deletion (Dilova et al. 2002).

Knowledge on metabolism and its regulation systems could be useful for improving the production of metabolites of interest (Cambon et al. 2006a). Efforts are being done to reduce ethanol content, due to health concerns, but also because global warming leads to an increase in sugar levels when grape berries reach maturity, leading to undesirable unbalances, with high ethanol and low acidity (Jones et al. 2005). Diverting the glycolytic flux from ethanol to glycerol is one of the best approaches to reduce alcoholic levels. Glycerol is involved in maintaining redox balance and is an osmolyte that prevent hyperosmotic shock. Besides, glycerol is a metabolite that have been reported to gives body, unctuosity and helps the perception of other aromas in wine, and it can be also positive in beermaking (Zhao et al. 2015). Glycerol is produced by diverting some of the trioses from glycolysis (dihydroxyacetone-P) into glycerol-3P by the action of glycerol-3P dehydrogenase. That process consumes NADH, so the net output is a reduction of fermentative flux and eventually a reduction of ethanol. Overexpression of *GPD1*, the main enzyme that produces glycerol achieves the expected phenotype, but due to redox unbalances, such modification leads to an increase of acetic acid, and that is not a desirable trait in fermented beverages (Remize et al. 1999). Additional deletion of acetic acid producing enzymes is therefore required (Cambon et al. 2006b). In this work, a deletion of RR repressor *MKS1* happens to increase glycerol without further increase in acetic acid in winemaking conditions. As a direct consequence, ethanol is also reduced. Similar output in glycerol production is seen in other strains of biotechnological interest, like brewing and baking yeast, so this seem to be a conserved regulatory mechanism that could be manipulated in order to modulate the amount of carbon fluxes during industrial fermentations.

## Materials and Methods

### Yeast strains and laboratory growth media

Haploid wine strain C9 was a gift by M. Walker (Walker et al. 2003). Wine commercial strains EC1118, T73, 71B and M2 comes from Lallemand Co. (Canada), brewing yeast SafAle US-05 is from Fermentis (Lesaffre, France) and baking yeast Cinta Roja from AbMauri (Spain). The whole *MKS1* gene was deleted from C9 by homologous recombination using the *kan*MX marker, which was amplified by PCR from the pUG6 plasmid using oligonucleotides with 40 bp homology to flanking regions (Guldener et al. 1996). The complete *RTG2* ORF was deleted from T73 strain by the same procedure. *MKS1* CRISPR-Cas9 deletions of diploid strains were made using plasmid pRCC-K, with the *<u>kan</u>MX* marker, which was a gift from Eckhard Boles (Addgene plasmid # 81191) (Generoso et al. 2016). Gene deletions were checked by PCR. Yeast transformations were done by the lithium acetate method (Gietz and Woods 2002).

Yeasts were propagated in rich YPD medium (1% yeast extract, 2% bactopeptone, 2% glucose). YPS and YPM changed the carbon source by 2% sucrose and 2% maltose, respectively. Solid plates contained 2% agar, and 20 μg/ml of geneticin for the selection of *kanMX* transformants. Minimal medium SD contained 0.17% yeast nitrogen base, 0.5% ammonium sulfate and 2% glucose (Adams et al., 1998). For the growth spot analysis, serial dilutions form stationary cultures in YPD were made and 5 μl drops were placed on selective media. The growth curves were obtained on a 96-well plate with 200 ml of each media inoculated a OD_600_ of 0.1 from a stationary culture in YPD. Plates were read with a Multiskan plate reader with shaking.

### Fermentations

C9*mks1*Δ standardized fermentations were carried out in a Sauvignon Blanc grape must by the method described before (Peltier et al. 2018). 20 mL screwed vials were tightly closed with screw cap with silicone/PTFE stoppers were filled with 11.5 mL of grape must inoculated with 2×10^6^ cell/mL and hypodermic needles were inserted for CO_2_ release. The fermentation temperature was maintained at 24 °C by an incubator. Vials were kept static or shaken at 175 rpm throughout fermentation using an orbital shaker. The fermentation kinetics was estimated by manually monitoring (2-3 times/day) the weight loss caused by CO_2_ release using a precision balance. The amount of CO_2_ produced according to time was modeled to estimate kinetics parameters: the maximal amount of CO_2_ released (CO2max in g.L^−1^), the lag phase (lp in h), the time to release 35, 50% and 80% of maximal expected CO_2_ after subtracting lp (t35-lp, t50-lp and t80-lp in h) and the average hexose consumption rate between 50% and 80% of CO2max (V50_80 in g.L^−1^.h^−1^). The concentrations of the following organic metabolites were measured at the end of fermentation: acetic acid, glycerol, malic acid, pyruvate and total SO_2_ using the respective enzymatic kits: (Megazyme, Bray, Ireland) following the manufacturer’s instructions. Glucose and fructose were assayed enzymatically (Stitt et al. 1989).

CRISPR-Cas9 edited commercial wine strains were grown in bobal red grape juice (Bodegas Murviedro, Requena, Spain) in conical tubes with 30 ml of juice at 24 °C (Orozco et al. 2012). CFU was followed by plating on YPD plates serial dilutions and counting colonies. Reducing sugars were measured with DNS (dinitro-3,5-salycilic acid) according to Miller’s method (Robyt and Whelan 1972). Other metabolites were measured with commercial kits (Megazyme Ltd, Bray, Ireland).

For brewing fermentations, a wort-like medium consisted of 180 g/l granulated malt supplemented with 10 g/l yeast extract was used (Garcia Sanchez et al. 2012). 11.5 ml or wort were used to fill screw-cap vials like the ones explained above at 25°C at 100 rpm and fermentation was followed by weight loss. For baking yeast a model liquid dough was used (Panadero et al. 2005) inoculated at 30 mg (dry weight) per ml, and fermentation was done in Erlenmeyer flasks at 30°C with low shaking (80 rpm).

## Results

### *MKS1* deletion increases glycerol and reduces acetic acid during winemaking

A phenomic analysis of nutrient signaling pathways by analyzing multiple deletions on haploid wine strain C9 in standardized microfermentations in white grape juice was done previously (Vallejo et al. 2020b). Mutation in RR repressor *MKS1* was performed and tested in these conditions, comparing static conditions (NS; not shaking) and shaking conditions (S), to analyze the effect of aeration. Evolution of fermentation was followed by weight loss (Figure 1A). The profile of fermentation is similar between both conditions, with a slightly lower fermentation speed for the *mks1*Δ strain. This effect is more evident from the beginning in the non-shaking conditions. For instance, time to reach 50% of CO_2_ production minus lag phase is always higher (Fig 1B). Metabolites of enological interest were measured at the end of fermentation, and *MKS1* deletion led to an increase of final glycerol (Fig. 1C) and a reduction of acetic acid (Fig. 1D) in both conditions. The glycerol produced by the mutant strain was even higher under shaking, but even in this condition that promotes the synthesis of acetic acid, the volatile acidity was low. Interaction between mutation and conditions was estimated, and lag phase time was the parameter that showed the biggest contribution of strain/shaking combination to the whole variance, 37% (Figure 1E). The start of fermentation is delayed in static conditions, but is reduced it in shaking conditions, giving indication that this pathway is responsive to oxygenation and influences the start of growth. Figure 1F shows the PCA plot that shows that mutants group together, and the correlation circle (Figure 1G) indicates that their influence in glycerol production is contributing to this fact.

**Fig 1.**
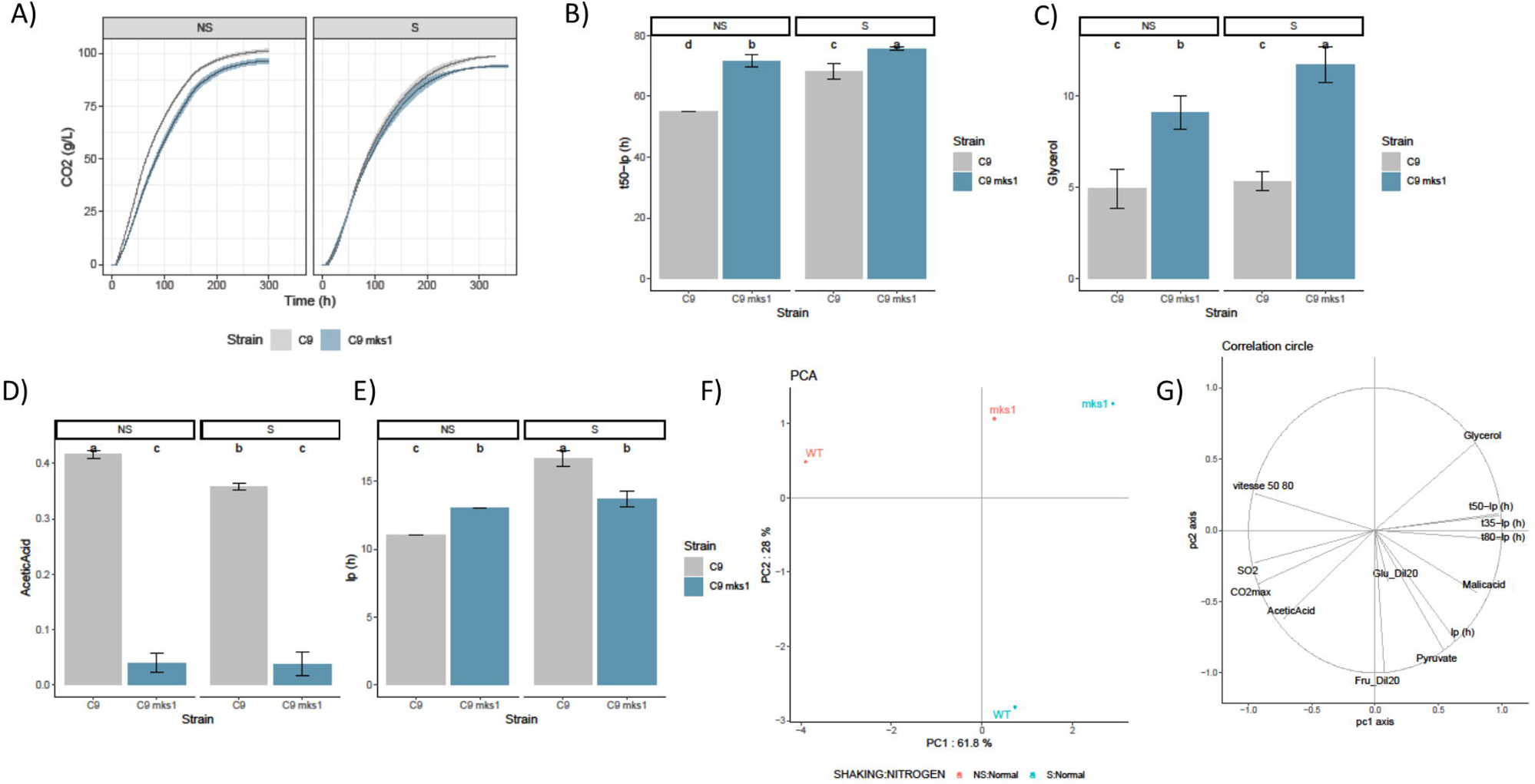
*MKS1* deletion impacts yeast metabolite production by an haploid wine yeast in microfermentations in white grape juice. A) Fermentations kinetic of strains C9 and C9 *mks1*Δ followed as CO_2_ production in static (NS) and shaking (S) conditions. B) Time to reach 50% of CO_2_ production minus lag phase. C) Glycerol production at the end of fermentation. D) Acetic acid production at the end of fermentation. E) lag phase. F) PCA analysis of kinetic and metabolic parameters of both strains in the two conditions. G) Correlation circle.

### Role of *MKS1* gene in commercial wine strains

Due to the interesting phenotype caused by *MKS1* deletion, it was tested in different conditions. First, instead of a haploid derivative, *MKS1* was deleted from a commercial diploid strain, EC1118. That was done by developing a CRISPR-Cas9 approach that enable us to delete both copies with a single transformation. Done that, microvinifications were carried out in natural grape juice (red must was used this time to test additional growth conditions) in open filled-in conical tubes. That allows to take samples along the process to measure sugar consumption and to analyze cell viability by plate count (Figure 2). Fermentation progress was measured by monitoring the reducing sugar consumption (Figure 2A). Again, in the mutant strain was slower in terms of fermentation speed, although it eventually reached completion. Regarding cell viability, the mutation caused a delay in growth and a lower maximum cell count (Figure 2B), suggesting that the slower fermentation is due to a lower cell number, not a lower fermentative power per cell. After reaching its maximum, cell viability decreased in both strains, so no big differences in lifespan were observed. Metabolites were measured at the end of fermentation, and again the mutation caused a significant increase in glycerol (Figure 2C), with a lower level of acetic acid (Figure 2D). The initial fermentations with C9 strain did not allow a standardized measurement of ethanol, but in this case, EC1118 *mks1*Δ produced a significant reduction in the final ethanol level, a reduction of 1g/100ml, a 9% less than produced by parental strain EC1118 (Figure 2E). Therefore, it seems there is a diversion in the glycolytic flux from ethanol to glycerol caused by this gene deletion. As this is a very interesting phenotype with biotechnological potential, the same CRISPR-Cas9 construct was used to delete the existing copies of *MKS1* gene in three other commercial strains: T773, 71B and M2. Fermentations were carried out in red grape juice as previously (Supplementary Figure S1). Strains had differences in the speed of fermentation measured by reducing sugars consumption (Supplementary Figure S1A), but the *MKS1* mutants always showed a slower fermentation, although all of them reach completion. Again, final ethanol production was significantly reduced (Supplementary Figure S1B), by 9% for the T73 and 71B genetic backgrounds, 10% for M2. Regarding glycerol (Supplementary Figure S1C), a doubling of the concentration of the metabolite was observed. Acetic acid (Supplementary Figure S1D) decreased in all cases, particularly in M2 genetic background. Therefore, the role of *MKS1* is consistent in all the genetic backgrounds tested and compatible with glycerol overproduction at expense of ethanol yield.

**Fig 2.**
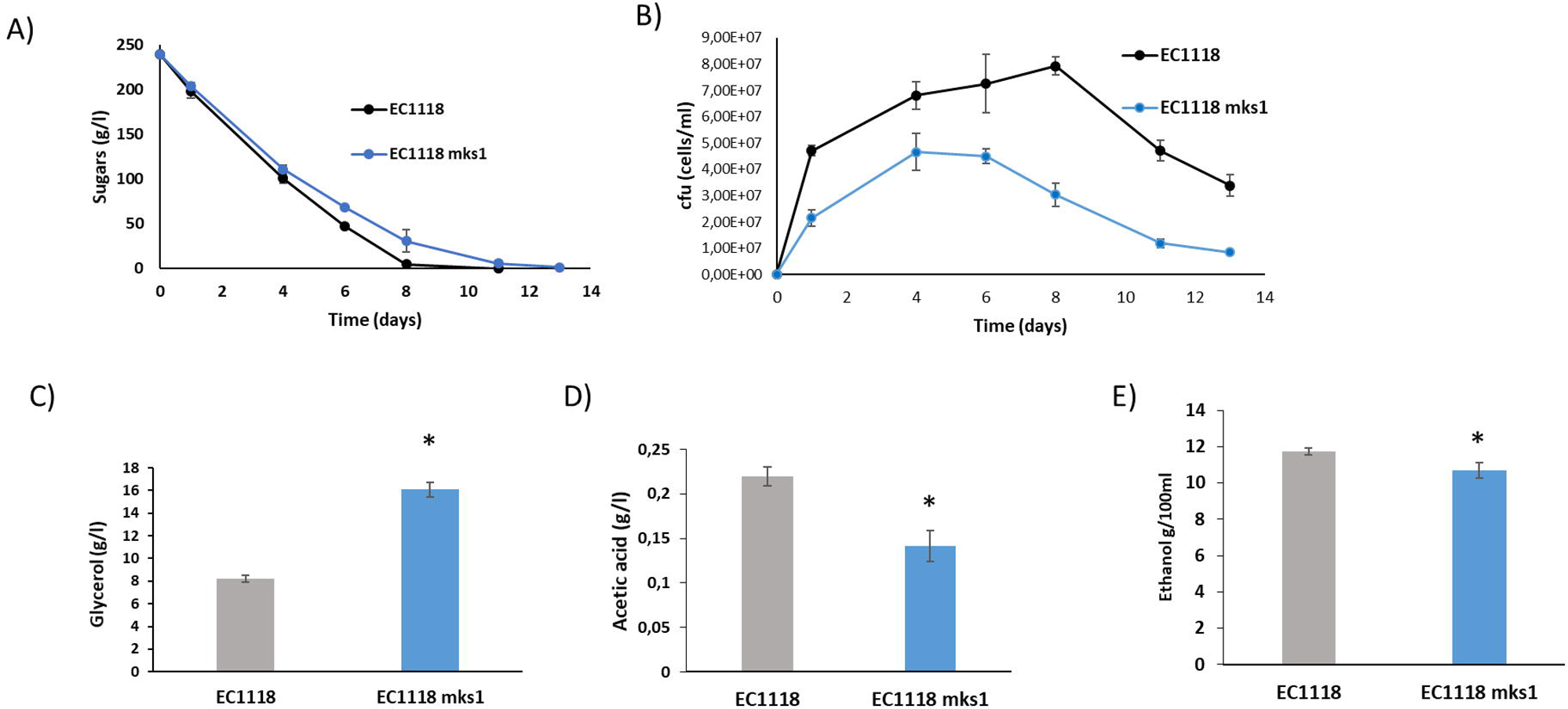
A CRISPR-Cas9 edited commercial wine yeast lacking *MKS1* increases glycerol and decreases ethanol production. EC1118 and EC1118 *mks1*Δ were grown in red grape juice. A) Total sugar consumption. B) Cell viability measured as CFU/ml. C) Glycerol production at the end of fermentation. D) Acetic acid production. E) Ethanol production. The shown values are the means of three replicates. Error bars represent standard deviation (* p≤0.05, two-tailed t-test).

In order to test the role of oxygen in the production of those metabolites of enological interest, a simple approach was done by comparing a static fermentation with a shaking condition on Erlenmeyer flasks where oxygenation was complete and full respiratory metabolism was allowed (Figure 3). M2 strain and its *MKS1* null mutant derivative were used as an example, as it is the strain with bigger ethanol reduction. In the shaking condition, reducing sugars were metabolized very fast compared to the non-shaking condition (Figure 3A). Again, the mutant strain showed a delay in the consumption of sugars (Figure 3A). As expected, the fully aerobic condition led to lower levels of ethanol, but again, the mutant strain had a decreased ethanol production compared to the wild type strain (Figure 3B). Similar levels of glycerol are produced by the parental strain M2 independently of shaking, and again *MKS1* deletion mutant increase this product in this condition, but to a lower level in the shaking condition (Figure 3C), maybe due to the diversion of reducing power to mitochondrial functions. As expected, aeration caused a high acetic acid production for wild type strain M2 (Figure 3D). However, this was reduced in M2 *mks1*, even to lower levels for this mutant in static conditions, causing a bigger overall reduction over five-fold. Retrograde response control D-lactic acid production, and that aspect was also measured (Figure 3E). In this case shaking decreases strongly lactic acid for the wild type strain, but in both conditions the absence of the *MKS1* gene led to an increase of this organic acid. That was not a common feature in all genetic backgrounds, as for instance it was seen in 71B strain but not in T73 strain in static conditions (data not shown).

**Fig. 3.**
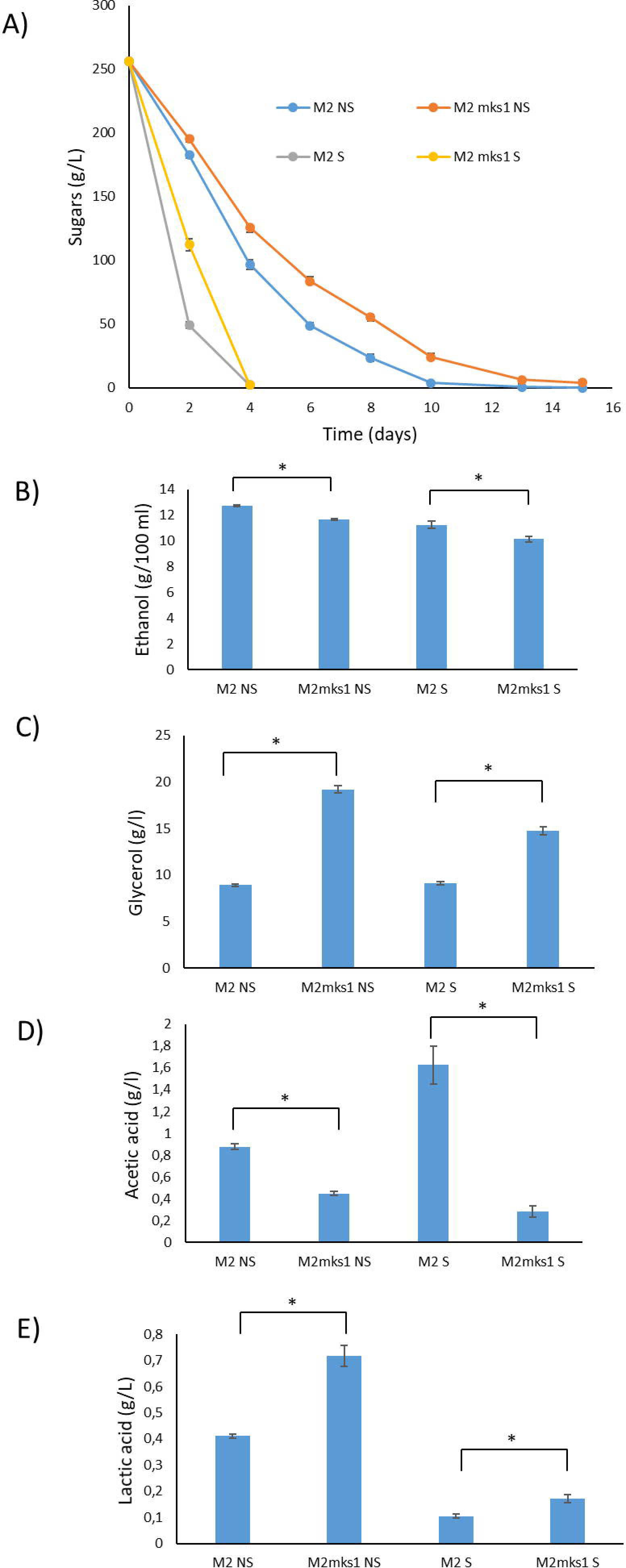
Full aeration in the presence of *MKS1* deletion does not produce high acetic acid levels. M2 and M2 *mks1*Δ were grown in red grape juice in flasks that were kept static (NS) or vigorously shaken (S). A) Reducing sugars consumption along fermentation. B) Ethanol production at the end of fermentation. C) Glycerol production. D) Acetic acid production. E) Lactic acid production. The shown values are the means of three replicates. Error bars represent standard deviation (* p≤0.05, two-tailed t-test).

### *MKS1* role in brewing and baking yeasts

As *MKS1* deletion has an impact in relevant metabolites during winemaking, it may affect the performance of other yeasts during the manufacturing of fermented foods. To test that, *MKS1* was deleted from a brewing yeast, Safale US-05 (Figure 4). To test its influence in malt fermentation, microvinifications in small vials used for standardized vinifications (see Figure 1) were used, and the advance of fermentation was followed by weight loss measurement (Figure 4A). *MKS1* deletion cause a delay in the start of brewing, and its overall profile is quite different. Mutant strain did not reach the same levels than the parental strain. Ethanol measured at day 8 (Figure 4B) indicated that in fact the mutant strain produces less ethanol than the parental strain. Glycerol (Figure 4C) and acetic acid (Figure 4D) were also measured. As happened in winemaking, there is a great increase in glycerol, so it seems that there was also a diversion from the glycolytic flux from ethanol to glycerol. Acetic acid was also reduced, as it happened in grape juice fermentation, so *MKS1* impact in metabolism seems to be similar independently of the genetic background and fermentation substrate used.

**Fig. 4.**
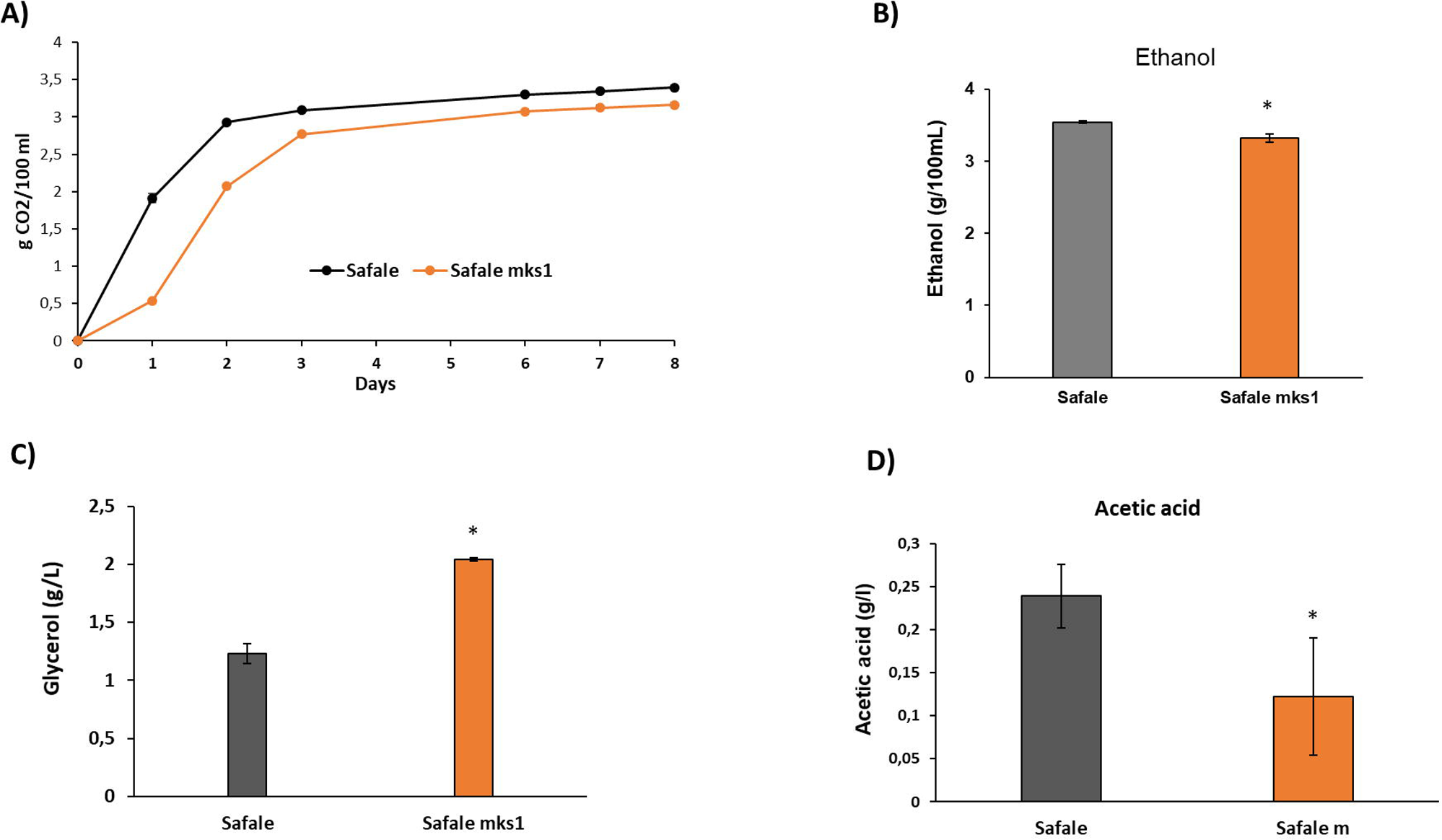
*MKS1* deletion produce high glycerol levels in brewing yeasts. SafAle US-05 and its M2 *mks1*Δ derivative were grown malt. A) Weight loss along fermentation. B) Ethanol production at the end of fermentation. C) Glycerol production. D) Acetic acid production. The shown values are the means of three replicates. Error bars represent standard deviation (* p≤0.05, two-tailed t-test).

The deletion in the baking strain Cinta Roja of *MKS1* gene was tested in a media called Liquid Dough that mimics a sweet dough (Panadero et al. 2005). That allows an easy inoculation and collection of samples. As breadmaking does not involve long times and a complete exhaustion of sugars is not expected, samples were taken over a period of 4 hours after inoculation and glycerol was measured as a marker of *MKS1* deletion impact on fermentation (Figure 5A). Glycerol was accumulated in the medium as fermentation progressed in the wild type, as expected. The increase in the *mks1*Δ strain is bigger, indicating that this mutation also increased glycerol in a laboratory-scale baking condition. Glycerol is produced to keep redox balance, but it is also a compatible osmolyte that prevents hyperosmotic stress, a feature that is important in a high osmolarity environment like dough. The *MKS1* deletion in all three industrial strains were tested in a spot test on plates with a high osmolarity (1.5 M KCl, Figure 5B). In the wine strain T73*, MKS1* deletion does not cause any improvement of stress tolerance, the colonies are even slightly smaller under stress. The negative effect is even more evident in the baking strain Cinta Roja. The *MKS1* deletion is more deleterious in the brewing strain SafAle, causing a growth defect even under non-stressful conditions, but in any case, the *MKS1* did not cause any relative improvement under hyperosmotic shock. Therefore, RR activation by *MKS1* deletion does not cause an improvement in osmotic stress tolerance in standard laboratory conditions.

**Fig 5.**
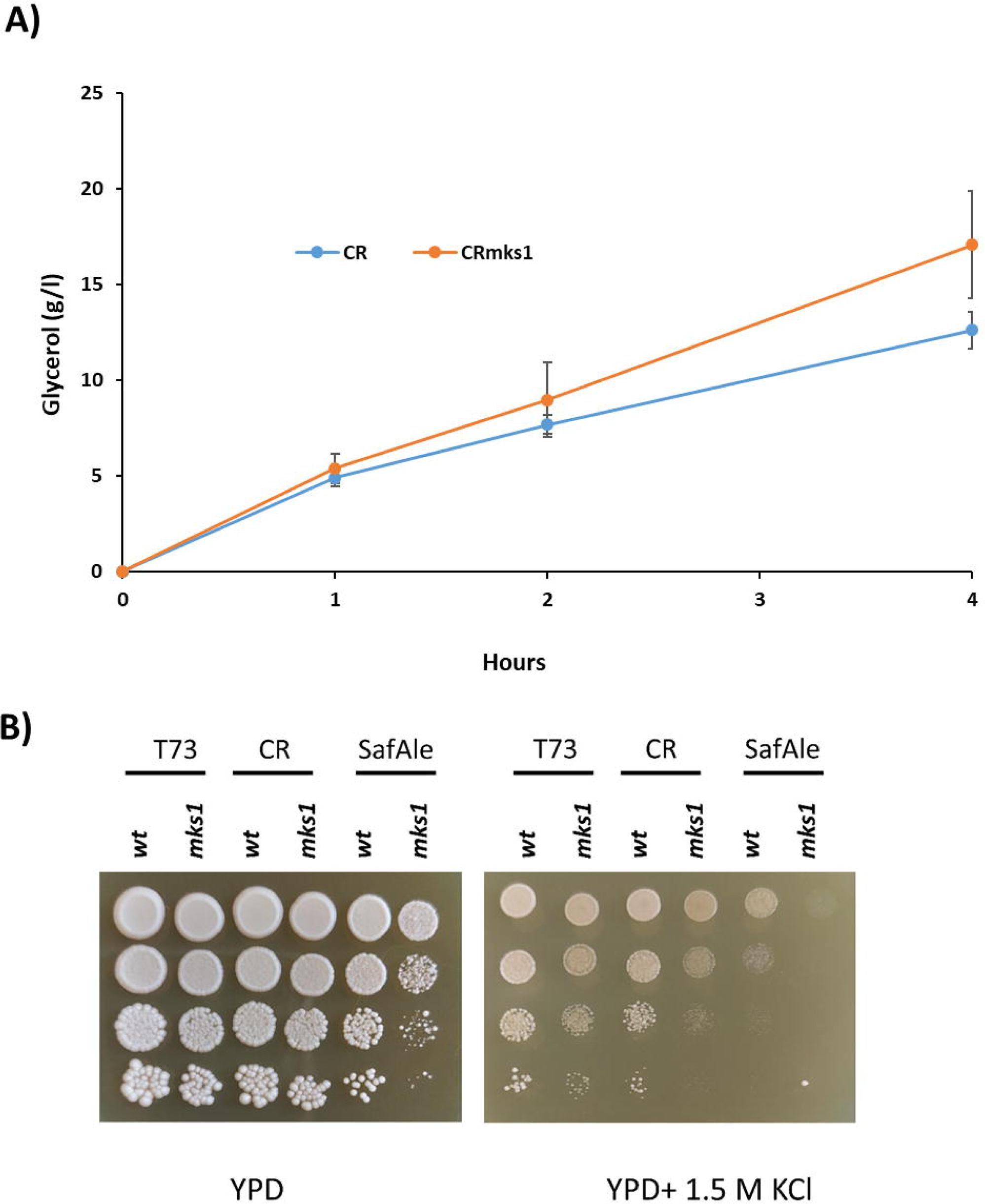
*MKS1* increase glycerol during baking. A) Glycerol production in a Liquid Dough medium by CintaRoja and its *mks1*Δ derivative. The shown values are the means of three replicates. Error bars represent standard deviation. B) Hyperosmotic tolerance in plates with 1.5 M KCl of deletion mutants of *MKS1* in wine (T73), baking (CR) and brewing (SafAle) yeasts.

### Role of *RTG2* in in wine yeasts

Due to the relevance in yeast performance showed by *MKS1* deletion, a mutation in the other key component of the RR pathway, the deletion of activator *RTG2* was performed in a wine yeast. Fermentation was followed in microvinifications like the ones depicted in Figure 1 and 4 by measuring weight loss (Figure 6A). There was a slight delay in fermentation in the mutant strain, but it reached completion (no reducing sugars were detected at end-time, data not shown). Therefore, there was not an opposite effect to *MKS1* deletion, indicating that balance RR is required for optimal yeast performance. There were no significant variations in ethanol (Figure 6B) nor acetic acid (Figure 6C). There was a very small but significant decrease in glycerol production (Figure 6D), as expected for a *MKS1* antagonist, but not in the degree that *MKS1* had. Therefore, Mks1 may have a wider range of targets than Rtg2, and *RTG2* deletion is not useful to influence the production of metabolites of enological interest.

**Fig 6.**
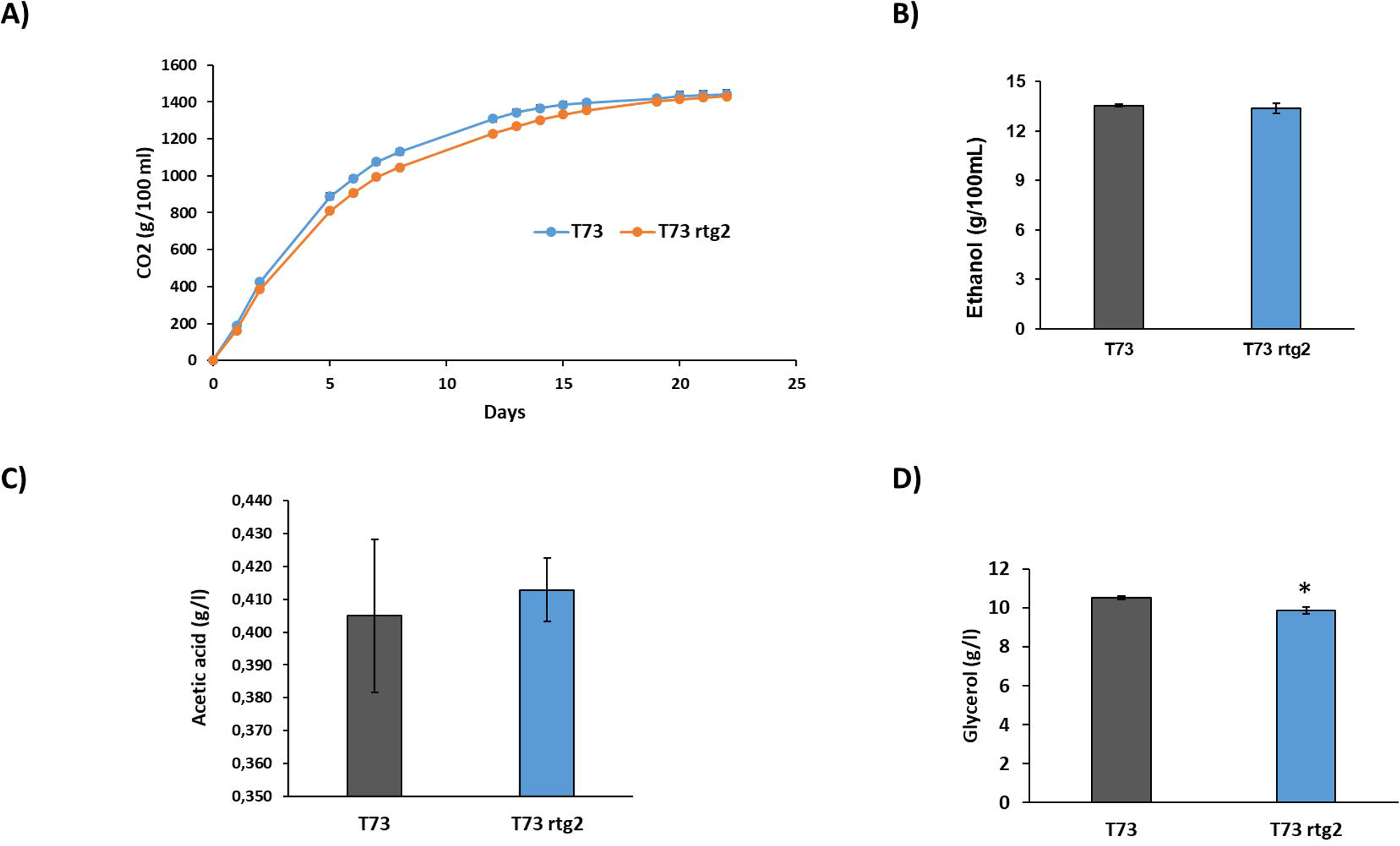
*RTG2* deletion has not a big impact on wine fermentation. T73 and T73 r*tg2*Δ strains were grown in red grape juice. A) Fermentation advance measured as weight loss. B) Ethanol production at the end of fermentation. C) Acetic acid production. D) Glycerol production. The shown values are the means of three replicates. Error bars represent standard deviation (* p≤0.05, two-tailed t-test).

### *MKS1* deletion impairs growth in a variety of carbon sources

As *MKS1* deletion mutant has a reduced fermentative power in several industrial media and reduced cell proliferation in grape juice fermentation, growth curves were performed in rich media with different carbon sources. The aim of this experiment is to determine if this an effect that is due to the harsh conditions of industrial growth media or an intrinsic condition of *MKS1* deletion regarding specific carbon source assimilation. First, the standard rich media YPD, that contains glucose as solo carbon source was used to test the deletions of *MKS1* and *RTG2* genes in wine strain T73 (Figure 7A). *rtg2*Δ strain showed a delay in growth, but eventually it reaches a similar saturation OD that the parental strain. However, *MKS1* deletion strain never reached a level of saturation and its growth speed is clearly lower. Therefore, Mks1 is required during exponential phase, but also to enter stationary phase. The role of *MKS1* was tested in two disaccharides of industrial relevance: sucrose, YPS (the main carbon source of molasses in which yeasts are propagated) and maltose, YPM (main carbon source in brewing and baking) (Figure 7B). Maltose is a worse carbon source than sucrose for this wine yeast, something not surprising considering that commercial wine yeast biomass is grown in molasses rich in sucrose. In both cases the pattern caused my *MKS1* deletion is similar: lower growth speed and lower maximum cell density for the mutant strain. Therefore, there is an intrinsic growth deficit in the *mks1*Δ strain regardless the carbon source. Deletion of *RTG2* has again a minor impact in these growth media (data not shown). The *MKS1* deletion on the brewing yeast SafAle was tested in two carbon sources YPD and YPM (Figure 7C). Maltose is relevant in the case of brewing and the parental strain is best adapted to grow in that medium than the wine yeast, although glucose is as expected the optimal carbon source. Again, *MKS1* mutation reduced growth speed and OD at saturation in both media, so the effect of *MKS1* on growth seems independent of the genetic background. Similar results were observed for the baking strain CintaRoja in similar conditions (Figure 7D).

**Fig. 7.**
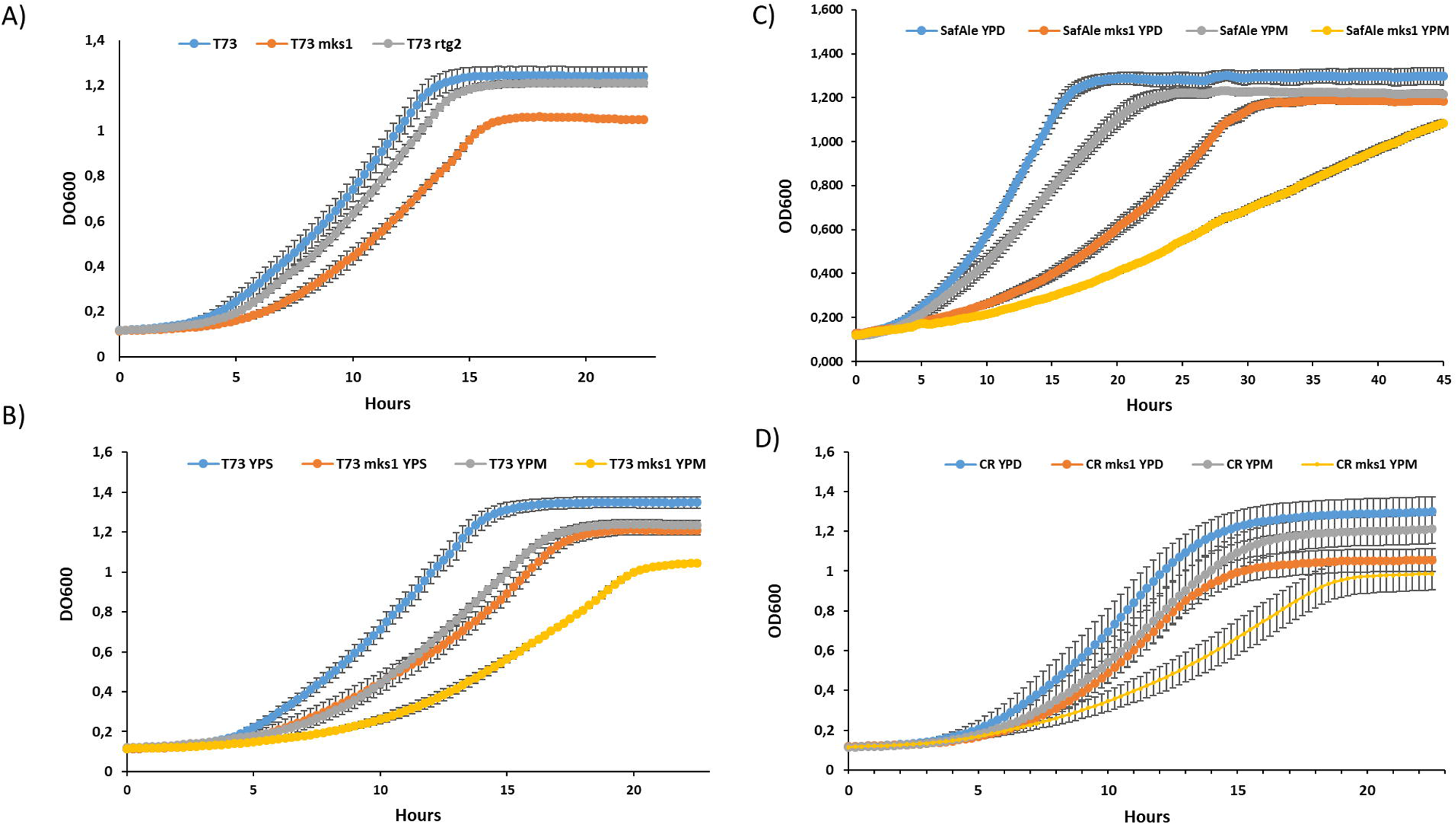
*MKS1* impacts growth in several carbon sources. Growth curves in 96-well microplate reader were obtained using rich media with different carbon sources: glucose (YPD), Sucrose (YPS) and maltose (YPM). A) Growth curves of wine yeast T73 and mutants T73 *mks1*Δ and T73 r*tg2*Δ in YPD. B) Growth of T73 and T73 *mks1*Δ strains in YPS and YPS. C) Growth of the *MKS1* deletion in the brewing yeast SafAle US-05 in YPD and YPM. D) Same as C for the CintaRoja baking strain. The shown values are the means of three replicates. Error bars represent standard deviation.

### *MKS1* deletion impacts amino acid biosynthesis

In order to probe the effect of *MKS1* deletion on amino acid metabolism, chemical inhibitors were used. Retrograde response increases the synthesis of α-ketoglutarate that is channeled to glutamate and lysine biosynthesis. For that reason *MKS1* was originally dubbed as *LYS80*, and its deletion increase tolerance to a lysine toxic analogue, 2-aminoethylcysteine (Feller et al. 1997). Mutations in *MKS1* and *RTG2* in wine strain were tested in spot analysis in the presence of this compound (Figure 8a). The phenotype is compatible with Retrograde Response regulation, and *MKS1* mutant is more tolerant to the compound and *RTG2* is more sensitive. The same pattern for *MKS1* deletion was seen in the SafAle brewing genetic background (Figure 8B) and in baking strain Cinta Roja (not shown). Therefore, the up-regulated synthesis of such mutant is conserved in all industrial yeasts. In order to test if this is a specific effect on some biosynthetic pathways and not a whole increase in amino acid biosynthesis, other toxic amino acid analogues were used (Figure 8A). 3-aminotriazole inhibits His3 and histidine biosynthesis, ethionine is a toxic analogue of methionine and canavanine of arginine. *MKS1* mutation does not increase the tolerance to these compounds, nor *RTG2* deletion decreases it significantly (*RTG2* deletion *per se* has a growth defect in minimal medium, so it may be targeting other pathways). Sulfometuron methyl that inhibits synthesis of branched amino acids valine and isoleucine in the mitochondria, and in this case, there is a growth defect in the *MKS1* deletion mutant, although *RTG2* deletion has a negligible effect. The toxic effect in *MKS1* was slightly relieved with the addition of valine and more so by the combination of valine+isoleucine, so it is targeting branch amino acids as expected, maybe by diverting precursors in the mitochondria to other amino acids.

**Fig. 8.**
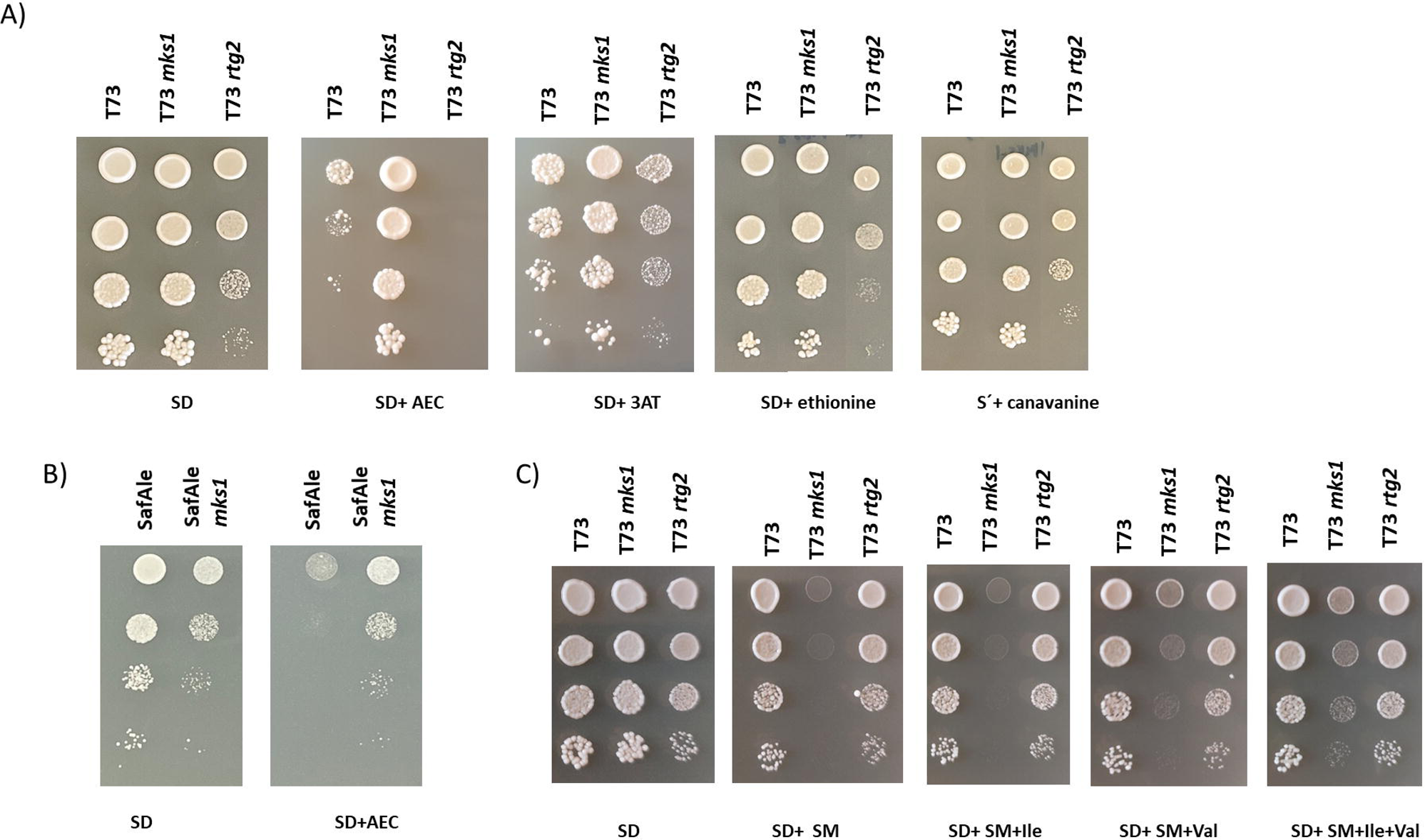
Mks1 plays a role in the biosynthesis of a subset of amino acids. Spot analysis of minimal medium plates using inhibitors of amino acid biosynthesis. A) Wine yeast T73 and mutants T73 *mks1*Δ and T73 r*tg2*Δ were tested against aminoethylcysteine (AEC), 3.aminotriazole (3AT), ethionine and canavanine. B) SafAle with *MKS1* deletion was tested for AEC tolerance. C) Wine yeast T73 and mutants T73 *mks1*Δ and T73 r*tg2*Δ were grown in the presence on sulfometuron methyl (SM) alone or in the presence of branched amino acids valine (Val) or isoleucine (Ile).

## Discussion

Nutrients are the guiding force behind the biotechnological processes that are carried out by microorganisms. A balanced growth medium guarantees the correct performance of the yeast *Saccharomyces cerevisiae* in its industrial tasks. Plant-derived substrates used for food production tend to be rich in carbon sources (mainly in the form of monosaccharides and disaccharides) but poor in nitrogen sources. Those industrial processes that aim at the full consumption of the carbon source, like winemaking, are more dependent on the presence of enough ammonium and amino acids to fulfill yeasts’ requirements. That is why sometimes addition of exogenous nitrogen source to the grape juice is required. Despite helping the progress of fermentation, this addition however can cause alteration in the aroma profile, that is largely dependent on amino acid metabolism (Gobert et al. 2019). Sugar molassesused for biomass propagation also require the addition of inorganic nitrogen to compensate for the excess of sucrose (Pérez-Torrado et al. 2015). Therefore, balance of nutrients is key for the success of any biotechnological process. Regulation of this processes is achieved through the nutrient signaling pathways (Conrad et al. 2014), and one of them is the Retrograde Response pathway, that channels carbon backbones from glycolysis to amino acid synthesis, so it is at the core of carbon/nitrogen relationship. We have focused in this study on the RR repressor, Mks1. Its deletion causes a decrease in wine fermentation speed in all genetic backgrounds and in both red and white grape juices, plus a defect on growth in different carbon sources in different industrial strains. The impact in fermentative performance of *MKS1* deletion was already detected in the screening of a wine yeast deletion collection (Peter et al. 2018). So, deregulation of the pathway *per se* is bad for yeast performance, probably because of resource misallocation. The mutation of RR activator *RTG2*, that should have the opposite effect, also impairs growth and fermentation, although to a lower stent. So, deregulation in this case could be worse than lack of activation. Biosynthesis of amino acids are therefore a very costly endeavor that has to be tightly regulated. The fact that *MKS1* deletion is sensitive to branched amino acid synthesis inhibitor sulfometuron methyl (Figure 8) may indicate that channeling resources to α-ketoglutarate derived amino acids cause deficits in other branches of nitrogen metabolism that may have a deleterious effect.

The most interesting result of this study is that fact that *MKS1* deletion alters carbon metabolism, leading to an increase of glycerol and decrease of ethanol in winemaking and brewing conditions. Glycerol overproduction at the expense of the glycolytic flux is a desired trait to get low alcohol wines, but overexpression of glycerol-producing enzymes leads to an undesirable increase in volatile acidity, probably due to redox-balance issues (Zhao et al. 2015) (Remize et al. 1999). Increasing the content of oxygen to induce a shift to a more respirofermentative metabolism is a way to decrease ethanol at the end of fermentation, but also has an increase of acetic acid production as a downside (Fig. 3; Tronchoni et al., 2022). In this deletion mutant acetic acid was not increased, in fact its levels decreased, so it shows a great potential for ethanol reduction with no organoleptic negative effect such as volatile acidity increase. *MKS1* deletion in industrial yeasts fits with the expected phenotypic trait of tolerance to lysine toxic analogue 2-aminoethylcysteine (Fig. 7). That is based in the fact that lysine is synthesized from α-ketoglutarate, and RR increase the synthesis of this precursor. RR also increases lactate dehydrogenase *DLD3* (Jazwinski 2013; da Cunha et al. 2015), and lactate is indeed increased in the mutant (Fig. 3). α-ketoglutarate comes from the condensation of oxaloacetate and acetyl-CoA by citrate synthase, and both the mitochondrial and peroxisomal isoenzymes are controlled by RR, so that may explain the reduction of acetate if acetyl-CoA is drained. The fact that the reduction is proportionally larger in conditions of aeration (Fig. 3) may indicate that the mitochondrial contribution may be more significant in this case. That depletion of acetyl-CoA in the cytosol may affect other processes, like lipid biosynthesis, impairing growth. Oxaloacetate comes from pyruvate in different pathways, including pyruvate carboxylase *PYC1*, another RR target. Any depletion from pyruvate will lead to decrease in ethanol and reducing equivalents could be channeled to glycerol production. Therefore, activation of RR fits with all phenotypic traits seen in different assays and backgrounds during fermentation of wine, beer and dough. Downregulation of RR by deletion of its activator *RTG2* should give an opposite effect, and it does mildly in one aspect, glycerol decrease, but it has not a consistent impact on ethanol. *MKS1* deletion has a well-known impact on canonical RR targets, but also it has a strong signature in the lysine biosynthesis (Dilova et al. 2002), while *RTG2* deletion does not show a control of this branch (Epstein et al. 2001). Therefore, *MKS1* and *RTG2* may have a different subset of targets, although both control RR in the production of α-ketoglutarate. *RTG2* deletion mutant sensitivity to 2-aminoethylcysteine (Fig. 8A) may be due to the decrease in lysine production by reduction of its precursor. In conclusion, manipulation of different aspects of RR may influence carbon metabolism, and increase of RR by *MKS1* deletions is a suitable way to produce low-alcohol beverages without increases in volatile acidity.

## Author Contribution Statement

VG, BV, EM, EP conducted experiments and analyze the data. PM and EM provide funding and review the manuscript. AA designed the experiments and wrote the manuscript.

## Data availability

All data generated or analysed during this study are included in this published article and its supplementary information files.

## Ethical approval

This article does not contain any studies with human participants performed by any of the authors.

