## Supplemental Figure 1 for "Up-regulation of Retrograde Response in yeast increases glycerol and reduces ethanol during wine fermentation"

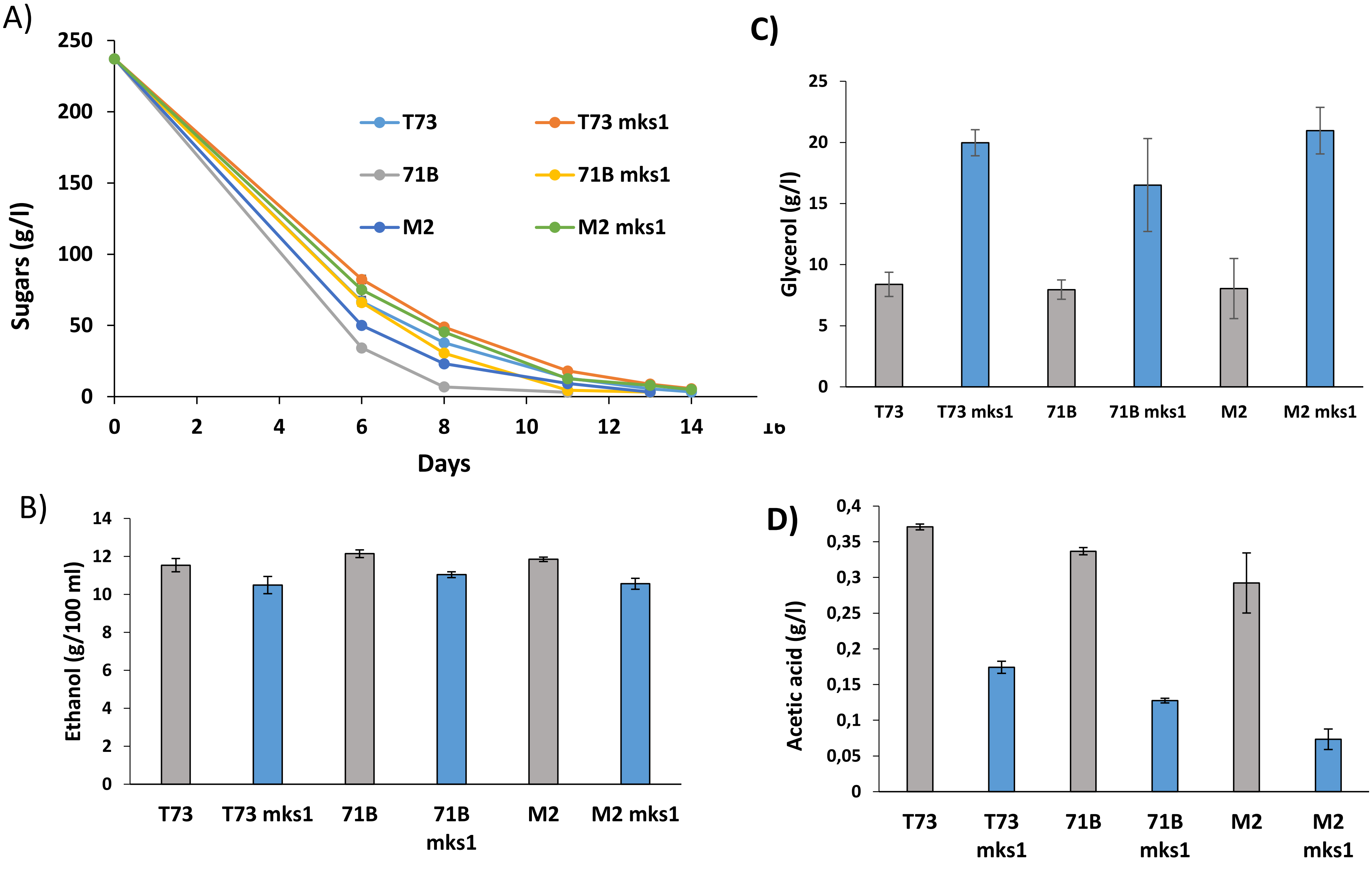


Supplementary Figure S1. CRISPR-Cas9 edited commercial wine yeasts lacking *MKS1* were grown in red grape juice. A) Total sugar consumption. B) Ethanol production at the end of fermentation. C) Glycerol production. D) Acetic acid production
